# Adrenergic Hypersensitivity Drives Ventricular Arrhythmias Following Loss of Plexin-Mediated Cardiac Innervation

**DOI:** 10.1101/2025.05.20.655085

**Authors:** Ching Zhu, Takako Makita, Emilio Y. Lucero, Arun Jyothidasan, Rhea Patel, Jessica J. Wang, Yang Cao, Howard A. Rockman, Kalyanam Shivkumar

**Author notes:** To whom correspondence should be addressed: Ching Zhu, MD, PhD, Duke Cardiovascular Research Center 213 Research Drive, Box 102156, Durham, NC 27710. **Conflict of interest statement**: Kalyanam Shivkumar: UCLA has patents developed by K.S. relating to cardiac neural diagnostics and therapeutics. K.S. is co-founder of NeuCures, Inc. All other authors: No conflicts of interest to declare.

## Abstract

**Background:** Ventricular arrhythmias (VAs) are a leading cause of death and arise from a combination of cardiac muscle injury and dysfunction of the intramyocardial sympathetic nerves that control cardiac electrophysiology. The adrenergic mechanisms by which intramyocardial nerves contribute to arrhythmogenesis are poorly understood. Semaphorin-plexin signaling pathways are responsible for developmental guidance of sympathetic nerves onto the heart and have previously been associated with VAs in humans.

**Objective:** To investigate adrenergic control of arrhythmogenesis, we probed the cardiac electrophysiology of a *Plexin-A3/-A4* double knockout mouse with loss of cardiac adrenergic nerves.

**Methods:** We studied cardiac structure and function using tissue clearing, immunohistochemistry, and echocardiography. We utilized ECG and optical mapping of action potentials to evaluate electrophysiologic responses to pharmacologic beta (β)-adrenergic stimulation and blockade. We measured circulating catecholamines and quantified β-adrenergic receptor (βAR) density in cardiac membranes. Finally, we performed a phenome-wide association study utilizing data from the UK Biobank to search for associations between *PLXNA4* and human arrhythmias.

**Results:** Mice with loss of plexin-dependent cardiac innervation had structurally normal hearts but displayed spontaneous VAs driven by adrenergic hypersensitivity, as well as increased cardiac βAR density. Several human *PLXNA4* variants were associated with arrhythmia phenotypes.

**Conclusion:** These data establish a model of VAs driven by enhanced adrenergic receptor signaling, in the absence of structural heart disease, which can be used to investigate adrenergic mechanisms of arrhythmogenesis and to identify novel antiarrhythmic targets.

## INTRODUCTION

Ventricular arrhythmias (VAs) result in sudden cardiac death, which accounts for one-fifth of mortality worldwide^1,2^. VAs arise from a combination of abnormal myocardial electrophysiology and dysfunction of the intramyocardial sympathetic/adrenergic nerves^3,4^. The adrenergic nerves within the heart direct the cardiac response to stressors and control every aspect of myocardial function, including arrhythmia susceptibility^5,6^. Modern anti-arrhythmic therapies rely on targeting cardiomyocyte electrophysiology as well as inhibiting adrenergic influence on the heart, but at present they have demonstrated limited efficacy in preventing VAs^7–9^. Both the excess (hyperinnervation) and the lack (hypoinnervation) of adrenergic axons within the ventricular myocardium have been associated with VAs^10–15^, but the mechanistic link leading from abnormal innervation to arrhythmogenic electrophysiology has not been elucidated.

Adrenergic axons infiltrate the heart during embryonic and early postnatal development, stimulated by nerve growth factors and spatially guided by trophic molecules released from coronary vessels^16–18^. This innervation process becomes quiescent in adulthood but is reactivated in acquired heart diseases such as myocardial infarction (MI) and heart failure^19–21^. One important mechanism directing cardiac innervation is semaphorin-3A signaling, which has previously been associated with cardiac arrhythmias in both humans and animal models^11,22,23^. Semaphorin-3a (*Sema3a*) knockout (KO) mice have abnormal cardiac innervation and arrhythmias^12^, but their arrhythmogenic mechanisms are challenging to study due to lethality in the early postnatal period^12,24^. The receptors Plexin-A3 and Plexin-A4, together with neuropilin co-receptors, mediate Sema3A signaling in developing sympathetic neurons, with Plexin-A4 transducing most of the Sema3A effect^25^. Mice deficient in Plexin-A4 have reduced cardiac innervation but are viable and fertile^25,26^, and their cardiac physiology has not been previously characterized.

In this study, we investigate the electrophysiologic consequences of abnormal cardiac adrenergic innervation in *Plxna3/a4* double KO (dKO) mice. Since the two plexins function redundantly^25,27^ in Sema3A signal transduction, we primarily studied the dKO to maximize the phenotypic effect of losing Sema3A signaling in cardiac innervation. We assess adrenergic axon density and cardiac structure using tissue clearing and immunofluorescence (IF) imaging in intact hearts. Using ECG recording *in vivo* and *ex vivo* optical mapping, we evaluated post-synaptic adrenergic responses in *Plxna3/a4* dKO mouse hearts and identify a phenotype of hyperadrenergic arrhythmogenesis.

## METHODS

### Animals

*Plxna3* and *plxna4* (MMRRC_030406-MU) alleles have been described previously^25^. All experiments complied with National Institutes of Health guidelines and were reviewed and approved by the UCLA Animal Research Committee, and both the MUSC and Duke Institutional Animal Care and Use Committees.

### Immunofluorescence and tissue clearing

Mice were given intraperitoneal (i.p.) heparin (100U) and euthanized with isoflurane and cervical dislocation. Once all reflexes subsided, a midsternal incision was made and animals were transcardially perfused with 50 mL ice-cold 0.01 M PBS containing 100U heparin followed by 50 mL freshly prepared, ice-cold 4% paraformaldehyde (PFA; EMS, RT 15714) in PBS. Hearts were post-fixed in 4% PFA overnight at 4 °C, washed, and stored in PBS with 0.01% sodium azide. Langendorff-perfused hearts were immersion fixed in 4% PFA overnight at 4 °C at the end of the functional experiments.

Hearts were stained and cleared using a modified iDISCO protocol^28^ as previously described^14^. Antibodies for cleared hearts were anti-TH (EMD Millipore, AB1542, 1:200) and donkey anti-sheep Alexa Fluor 647 (Jackson ImmunoResearch, 713-605-147, 1:300) diluted in 0.01 M PBS with 0.2% Tween-20 and 10 mg/ml heparin (PTwH). Primary and secondary Ab were replenished approximately halfway through the incubation period. Hearts were stored in benzyl ether (Millipore Sigma, 108014 ALDRICH; refractive index: 1.55) for up to 7 days prior to imaging.

For qualitative assessment of lateral left ventricular (LV) myocardial and atrioventricular (AV) nodal sections, 300 μm-thick vibratome sections were created from fixed hearts and immunolabeled with anti-HCN4 (1:400; Alomone Labs APC-052), anti-TH (1:400: Millipore AB1542) and anti-SYP (1:500; SCBT sc-17750) antibodies overnight at 37°C, and followed by Alexa Fluor-conjugated secondary antibodies (Life Science) for 3 hr at 37°C. Immunolabeled sections were counterstained with DAPI and cleared with ScaleU2^29^.

### Confocal imaging and image analysis

Whole hearts were mounted in benzyl ether with adhesive plastic spacers (Sunjin Labs, IS012 and IS013). Images were acquired on a confocal laser scanning microscope (Zeiss, LSM 880) at 5x magnification. Whole heart images were taken in tiles with XY-resolution of 1.661μm and Z-resolution of 8.29μm. Heart sections were flat-mounted to slides and imaged at 40x magnification.

To quantify TH-positive axons in whole hearts, tiles were stitched with Zeiss Zen software and a maximum intensity projection (MIP) was created of each heart. The TH-positive pixel area was measured by binary threshold selection using Fiji-ImageJ software. This was divided by the whole heart area, as measured by muscle autofluorescence, to calculate percentages.

### Echocardiography

Mice were anesthetized with 3% isoflurane and maintained with 1% isoflurane in 95% oxygen. Anesthesia was adjusted to obtain a target heart rate of 470 ± 50 beats per minute (bpm). Transthoracic echocardiography was conducted with a Vevo 3100 high-frequency, high-resolution digital imaging system (VisualSonics) equipped with a MX550S MicroScan Transducer. Fur was removed a day prior to imaging studies. Mice were immobilized in a supine position at a 20-30 degree incline. Ultrasound gel was warmed to approximately body temperature and applied to the area overlying the heart. A parasternal short axis view, indicated by the presence of papillary muscles, was used to obtain M-mode images for analysis of fractional shortening, ejection fraction (LVEF), and other cardiac structural parameters. All parameters were measured two to three times, and means were used for analysis. At the end of the procedures, all mice recovered from anesthesia without difficulties.

### Electrocardiography and analysis

ECG data was recorded in awake, freely moving mice by implanting radio telemetry devices (ETA-F10, Data Science International). Mice were anesthetized with 4% isoflurane and maintained with 2% isoflurane. Telemetry units were implanted into the peritoneal cavity. The electrical leads of the telemetry units were tunneled through the abdominal wall with the free ends secured to the pectoral muscles using non-absorbable suture in a lead II orientation. Mice were recovered for 14-20 days following telemeter implantation surgery to ensure minimal lead motion artifact and to avoid the effects of postoperative pain and inflammation on baseline autonomic state. Mice were housed singly in cages over antenna receivers connected to a computer system with Ponemah software (Data Science International) for recording. Approximately 48 hours of continuous ECG recording were collected for each mouse, with brief interruptions for the intraperitoneal injection of isoproterenol (1 µg/g) and propranolol (4 µg/g).

Anesthetized ECGs were performed in mice during 2% isoflurane inhalation, using platinum needle electrodes (AD Instruments) placed subdermally in the lead II configuration. Recording was performed using a Powerlab 35 acquisition system (AD Instruments). Once stable baseline heart rate was confirmed for 5 minutes, isoproterenol 1 µg/g was administered by intraperitoneal injection and data was collected for an additional 20 minutes after isoproterenol administration. LabChart 8.0 software (AD instruments) was used for anesthetized ECG analyses.

Analysis of telemetry recordings, including premature ventricular contraction (PVC) and ventricular tachycardia (VT) quantification, was performed using ecgAUTO software (Emka Technologies). ECG interval measurements were taken from the average of a 5-min recording at 9am. Baseline heart rates (HR) were averages of 2 hours prior to isoproterenol administration, starting at 4pm. Post-drug arrhythmia quantification was performed for the 2 hours following isoproterenol or propranolol administration. The arrhythmia severity scores were defined as: 0 = no ventricular ectopy, 1 = isolated PVCs, 2 = bigeminy or couplets, 3 = VT (3 or more continuous ectopic beats).

### Optical Mapping of Action Potentials

Optical mapping of membrane voltage was performed as previously described^14^, during both sinus rhythm and ventricular ectopy induced by isoproterenol 100nM added to the Langendorff perfusate reservoir. Data were analyzed using BV Workbench (SciMedia) and ElectroMap^30^. Activation maps were created using the maximum second derivative of fluorescence (F) intensity over time (d^2^F/dt^2^).

### Langendorff perfusion and pseudo-ECG

Mice were injected with intraperitoneal heparin (100 U) 10-15 minutes prior to sacrifice per protocol by anesthesia with 5% isoflurane followed by cervical dislocation. Hearts were removed immediately and perfused via the aortic root with Tyrode’s solution (130mM NaCl, 1.25mM CaCl_2_, 5mM KCl, 1.2mM NaH_2_PO_4_, 1.1mM MgCl_2_, 22mM NaHCO_3_, and 50mM dextrose, aerated by bubbling 95% oxygen and 5% carbon dioxide via a fritted gas dispersion tube). Hearts were immobilized and immersed in Tyrode’s solution bath within a custom, 3-D printed chamber to reduce motion artifact. Perfusate and bath temperature were maintained at 36.6-37°C. Aortic root pressure was monitored with a Powerlab Bridge Amp (AD Instruments) and maintained constantly at 70-80 mmHg for the duration of the experiments, with meticulous flow rate adjustments (3-6 ml/min). Following confirmation of stable heart rate for 20 minutes, isoproterenol was administered via addition to the perfusate reservoir at 100nM concentration, and an additional 20 minutes of data was recorded.

Pseudo-ECG was recorded with platinum needle electrodes (Grass Instruments) stabilized in Sylgard 184 silicone elastomer (Electron Microscopy Sciences) using a P511 amplifier (Grass Instruments) connected to a Powerlab 35 acquisition system (AD Instruments). ECG analysis was performed using LabChart software (AD Instruments).

### Determination of plasma catecholamine levels

Mice were euthanized by isoflurane overdose and thoracotomy was performed. Blood was collected by puncture of the cardiac apex and aspiration through a 20G needle into a syringe primed with 0.5M EDTA. Blood samples were centrifuged at 2000 x *g* for 10 minutes at room temperature, and the plasma supernatant was stored immediately at –80°C for a maximum of 3 weeks prior to measurement of catecholamine levels. Plasma epinephrine and norepinephrine levels were measured using an enzyme-linked immunosorbent assay (ELISA) kit (Eagle Biosciences EA613/192). Data were analyzed using 4PL regression in Prism 10 (GraphPad).

### Membrane extraction from cardiac ventricular tissue

Mice were sacrificed per protocol by anesthesia with 5% isoflurane followed by cervical dislocation. Hearts were dissected, ensuring complete removal of atria, pulmonary and aortic vasculature, then flash frozen in liquid nitrogen and stored at –80°C until needed. Frozen ventricles were resuspended in ice-cold lysis buffer (Tris–HCl 25 mM, EDTA 5 mM, and 1X of the proteinase inhibitor mix aprotinin and leupeptin [LB]; pH 7.4), minced, and dounce homogenized (30 strokes). Homogenates were centrifuged at 500 g for 5 minutes at 4°C. The supernatant was discarded, and the pellet was resuspended in binding buffer (Tris–HCl 75 mM, EDTA 2 mM, MgCl2 12.5 mM, and 1X LB mix; pH 7.4). The homogenates were centrifuged at 35,000 g for 30 minutes at 4°C to obtain the membrane fraction. The supernatant was discarded, and the pellet was resuspended in binding buffer with 10% glycerin, homogenized, and stored at −80°C until needed.

### Radioligand Saturation Binding

Protein quantification of cardiac membrane fractions was performed using the bicinchoninic acid (BCA) protein assay following the manufacturer’s instructions. Then, 18 to 20 μg of protein diluted in 50 μl of binding buffer was added per well, followed by 50 μl of binding buffer or propranolol (30 μM) to determine non-specific binding (NSB). Finally, increasing concentrations of Iodo-(-)-Cyanopindolol, [125I] (125I-CYP) from 35.7 to 300 pM were used to measure total binding (TB) at a constant volume of 200 μl/well. The mixture was incubated for 90 minutes at room temperature to reach equilibrium. The radioactivity was measured as counts per minute using a gamma counter (PerkinElmer, USA). The saturation binding curves are reported as specific binding (SB), where SB = TB − NSB. The receptor density (Bmax) was determined using one site-specific binding regression curve (Graphpad Prism 10). The Bmax values were converted from counts per minute to fmol/mg of protein.

### Phenome-wide association study

To evaluate the association between plexins and arrhythmias in humans, we searched for associations between all phenotypes containing the word “arrhythmia” and the gene “*PLXNA4*” via the AstraZeneca PheWAS portal (https://azPheWAS.com/). This analysis was based on data from the UK Biobank, which included approximately 15,500 binary phenotypes and 1,500 continuous phenotypes derived from around 500,000 participants who had undergone exome sequencing^31^. The detailed methodology is documented in the original publication.

### Statistics

All statistical analyses for comparison are indicated in figure legends or main text and were performed in Prism 10 (GraphPad), unless otherwise noted in the text. Statistical tests were all two-tailed, unless otherwise noted in the text.

## RESULTS

### *Plxna3/a4* dKO mice exhibit cardiac adrenergic denervation but have structurally normal hearts

To evaluate cardiac adrenergic innervation in *Plxna3/a4* dKO mice, we performed whole-mount immunofluorescence (IF) staining for the adrenergic neuron marker tyrosine hydroxylase (TH) in intact hearts and cleared them using a modified iDISCO+ protocol^28^. Confocal imaging revealed a significant global reduction in cardiac adrenergic nerves in *Plxna3/a4* dKO hearts compared to wild-type (WT) controls (Figure 1A). Quantification of TH staining as percent of total heart area in maximum intensity projection (MIP) images showed that TH-positive axons were less than half as dense in *Plxna3/a4* dKO hearts compared with WT hearts (Figure 1B, Cliff’s δ=1, n=4 WT and 3 dKO). Though most of the heart area was ventricular in these quantitative analyses, stratification of the data also showed decreased atrial innervation (Supplemental Figure 2A-B). Qualitatively, denervation in *Plxna3/a4* mice was observed throughout the whole heart, as demonstrated by decreased staining for TH and the synaptic marker synaptophysin (SYP) in tissue sections from the working myocardium of the lateral LV and from the AV nodal region as defined by anti-HCN4 staining (Figure 1C-D).

**Figure 1.**
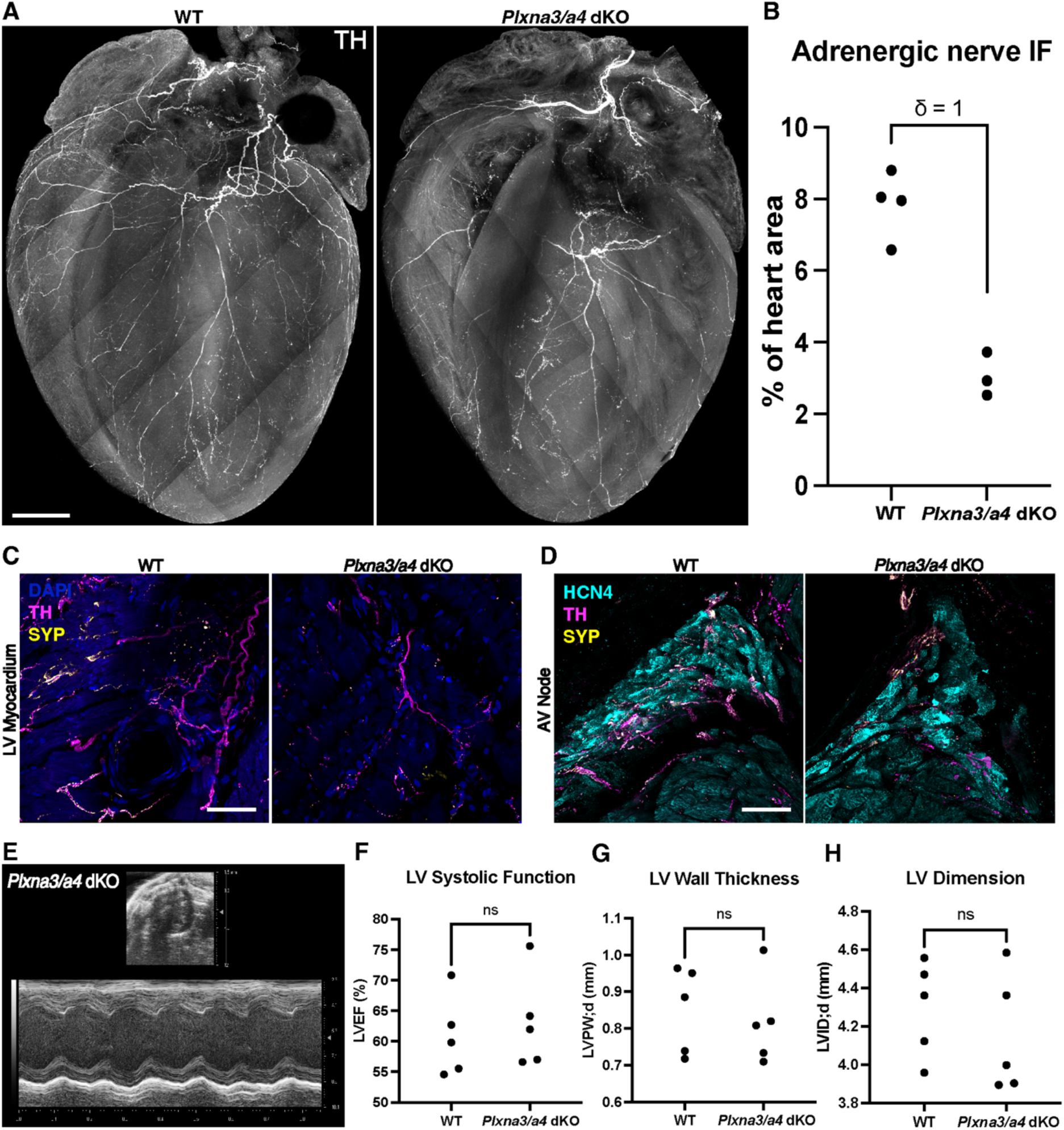
Decreased adrenergic innervation and normal cardiac structure in *Plxna3/a4* dKO hearts. **A**) Representative maximum intensity projection images of whole-mount IF in iDISCO-cleared hearts demonstrates globally reduced adrenergic nerve marker TH in *Plxna3/a4* dKO mouse ventricles (right image) compared with WT (left image). **B)** TH quantification, calculated as percent of whole heart area, confirms *Plxna3/a4* dKO hearts have less than half the adrenergic nerve IF compared with WT hearts (Cliff’s δ = 1, each dot represents one mouse). **C,D)** IF of tissue sections from *Plxna3/a4* dKO hearts qualitatively confirms the adrenergic denervation phenotype in both working lateral LV myocardium and conduction system (AV nodal region) defined by HCN4. **E-H)** Transthoracic echo demonstrates *Plxna3/a4* dKO mice have normal heart structure and LV systolic function compared with WT mice (Mann Whitney tests, p>0.05, each dot represents one mouse). SYP = synaptophysin, used to stain nerve endings. ns = not significant. Scale bar in **(A)** = 200μm, in **(C)** and **(D)** = 100µm.

Despite the significant reduction in adrenergic innervation, *Plxna3/a4* dKO mice showed no abnormalities in cardiac structure or systolic function. Transthoracic echocardiography demonstrated normal LV ejection fraction (LVEF), posterior wall thickness (LVPW;d) and end-diastolic dimension (LVID;d) in *Plxna3/a4* dKO mice compared to WT controls (Figures 1E-H, Mann-Whitney tests, p > 0.05 for all parameters, n = 5 mice per group). There was no fibrosis seen in tissue sections from *Plxna3/a4* dKO hearts stained with Masson’s trichrome (Supplemental Figure 1).

### *Plxna3/a4* dKO mice have spontaneous ventricular arrhythmias that are exacerbated by beta-adrenergic stimulation and attenuated by beta-adrenergic blockade

To determine whether adrenergic denervation affects underlying electrophysiology in *Plxna3/a4* dKO mice, we performed 48-hour continuous ECG recordings in conscious, freely-moving mice using implanted telemeters. Both WT and *Plxna3/a4* dKO mice had normal PR, QRS and QT intervals, and their resting HRs were not significantly different (Supplemental Figure 3A-B). While WT mice displayed normal sinus rhythm without any observed ventricular ectopy at baseline (Figure 2A), *Plxna3/a4* dKO mice frequently exhibited spontaneous premature ventricular contractions (PVCs) that were overall pleiomorphic in nature (Figure 2B-C). These PVCs occurred with both fixed (Figure 2B) and variable coupling intervals (Figure 2C). *Plxna3/a4* dKO mice also had episodic ventricular tachycardia (VT) (Figure 2D). Quantification of arrhythmia severity demonstrated significantly higher arrhythmia scores in *Plxna3/a4* dKO mice compared to WT controls at baseline (Figure 2E, one-tailed Mann-Whitney test, p = 0.0303), and the frequency of these arrhythmias was also increased (Figure 2F, one-tailed Mann-Whitney test, p = 0.0011). Premature atrial contractions (PACs) were also seen in the same telemetric ECG data, but there was no difference in PAC burden between *Plxna3/a4* dKO mice and WT controls (Supplemental Figure 2C-E).

**Figure 2.**
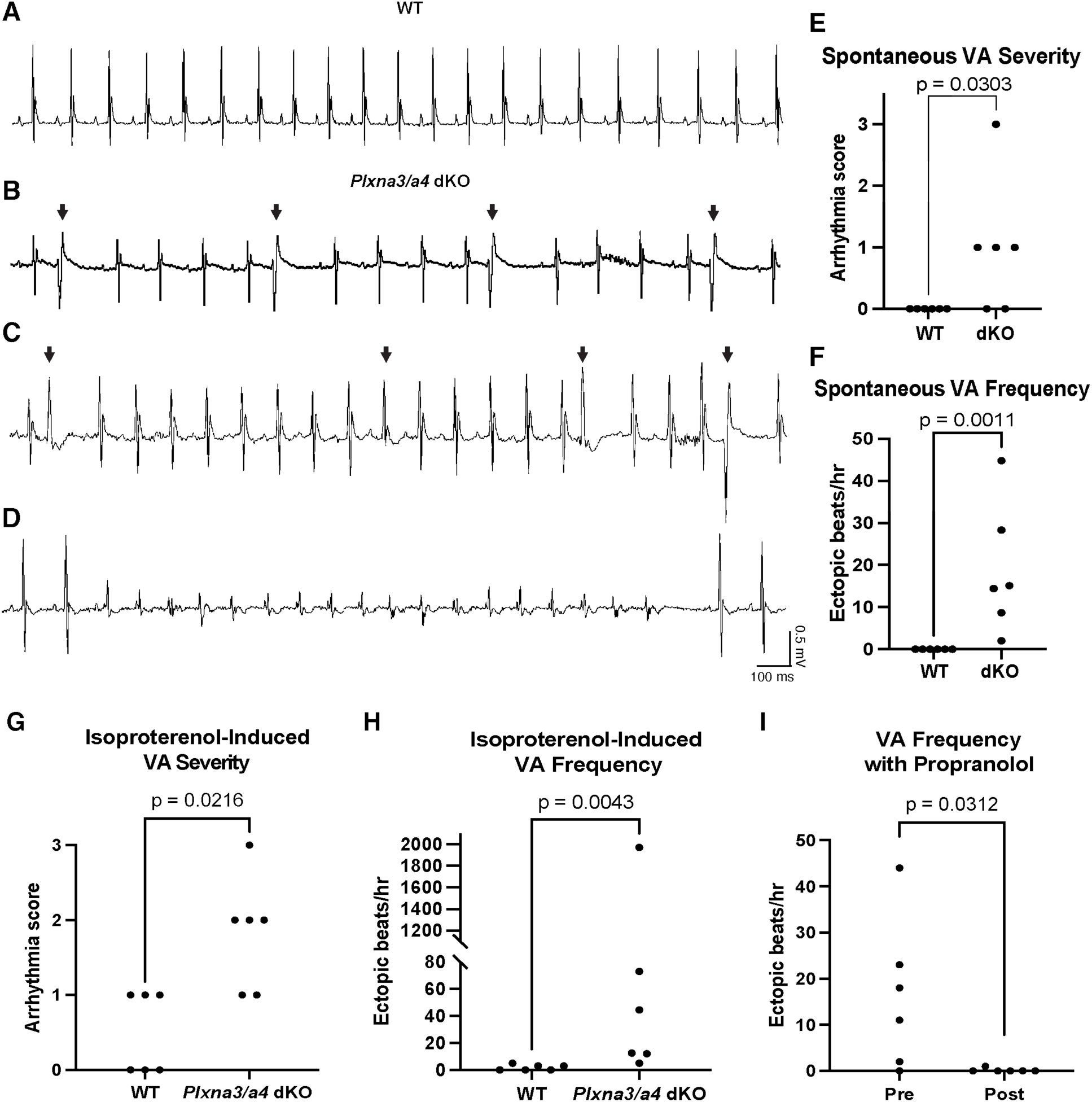
Spontaneous ventricular arrhythmias in *Plxna3/a4* dKO mice are exacerbated by beta-adrenergic stimulation and inhibited by beta-adrenergic blockade. **A**) Representative ECG recording from a conscious, freely moving WT mouse shows normal sinus rhythm. **B)** Frequent, spontaneous PVCs of fixed coupling interval in *Plxna3/a4* dKO mice. **C)** PVCs of variable coupling intervals with fusion beat (third arrow-demarcated beat) and **D)** VT in *Plxna3/a4* dKO mice. **E)** Ventricular arrhythmia (VA) severity and **F)** frequency in *Plxna3/a4* dKO mice versus WT. **G)** VA severity scores following isoproterenol (1 µg/g i.p.) injection. **H)** Frequency of induced VAs in the 2 hours following isoproterenol administration. **I)** VA frequency in *Plxna3/a4* dKO mice during 2-hour baseline recording (pre) compared to the 2 hours after (post) propranolol administration (4 µg/g i.p.) administration. P-values in **(E-F)** are from one-tailed Mann-Whitney tests, **(G-H)** from two-tailed Mann-Whitney tests, and in **(I)** is from one-tailed Wilcoxon paired rank test. Each dot represents one mouse. VA severity scores: 0 = no PVCs, 1 = isolated PVCs, 2 = couplets or bigeminy, 3 = VT (3 or more consecutive ventricular ectopic beats).

VA severity in *Plxna3/a4* dKO mice was amplified following intraperitoneal administration of the beta (β)-adrenergic agonist isoproterenol (1 μg/g) (Figure 2G, Mann-Whitney test, p =0.0216). The frequency of VAs, defined by ventricular ectopic beats per hour, was also significantly higher in *Plxna3/a4* dKO mice following isoproterenol administration (Figure 2H, Mann-Whitney test, p = 0.0077). The peak HR following isoproterenol administration was not changed in *Plxna3/a4* dKO mice compared to WT (Supplemental Figure 3B). Administration of the β-blocker propranolol (4 μg/g, i.p.) essentially abolished VAs in *Plxna3/a4* dKO mice (Figure 2I, one-tailed Wilcoxon paired rank test, p=0.0312) in the 2 hours after injection, when compared with the 2 hours prior to propranolol administration.

We confirmed the ventricular origin of the observed arrhythmias by performing optical mapping of action potentials (APs) in a representative *Plxna3/a4* dKO mouse heart in Langendorff (Figure 3). We imaged the ventral surface of the heart, including both the left atrium and ventricles. We induced VAs of two morphologies in the Langendorff preparation by perfusing isoproterenol 100 nM. Activation maps showed that while ventricular activation was earliest at the exits of the bundle branches in sinus rhythm (Figure 3B), it was earliest from the lateral LV during PVC #1 (Figure 3D) and from the lateral RV during PVC/VT #2 (Figure 3F). The relative isochronal crowding in VA activation maps compared to the rapid activation in sinus rhythm suggests lower velocity of impulse propagation – possibly due to origin from myocardium rather than conduction system. Simultaneous mapping of atrial and ventricular APs showed atrioventricular (AV) dissociation during VT (Figure 3G).

**Figure 3.**
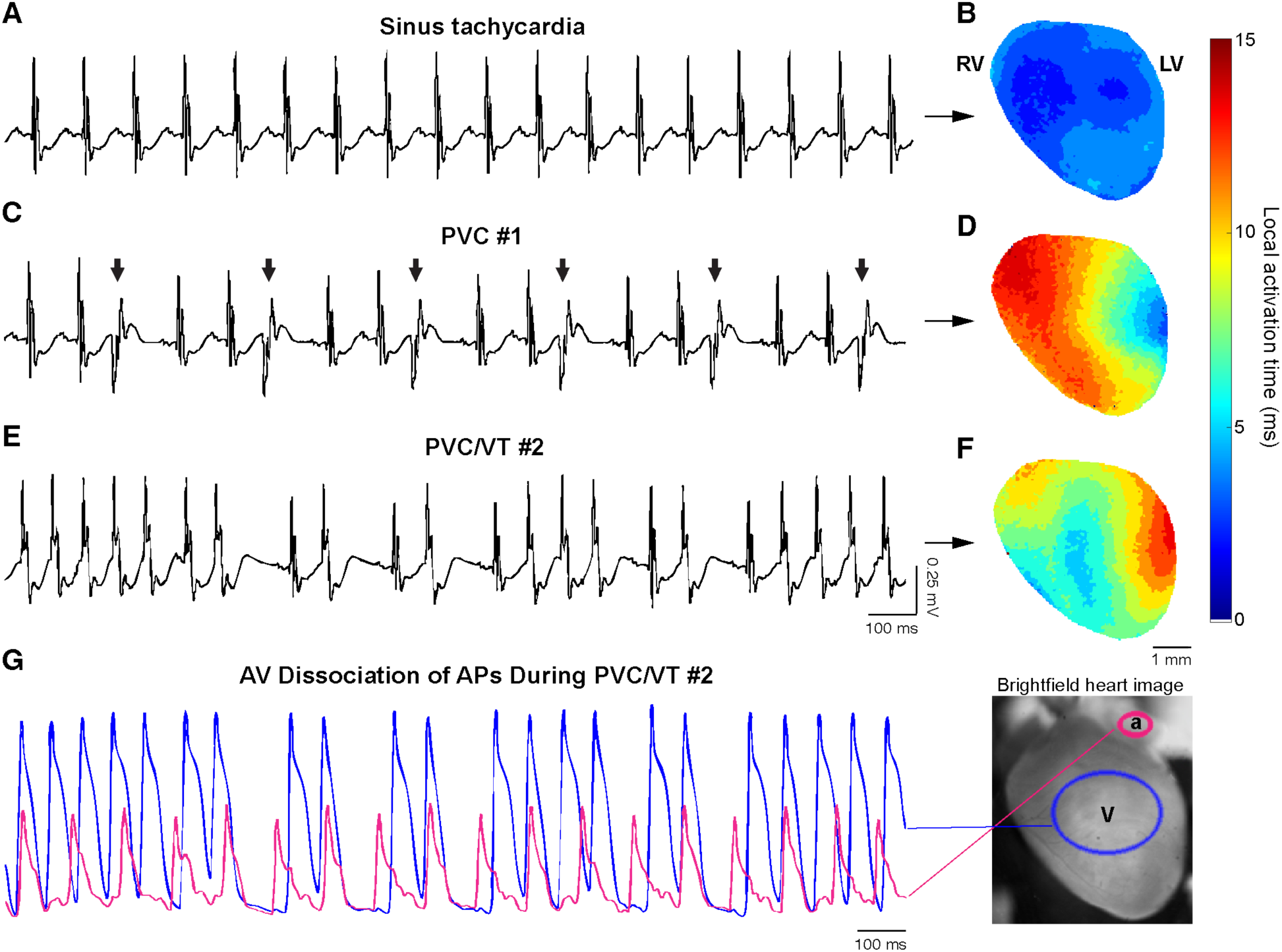
Optical mapping of action potentials during isoproterenol-induced ventricular arrhythmias in a representative *Plxna3/a4* dKO mouse heart. **A**) Representative pseudo-ECG recording from a Langendorff-perfused *Plxna3/a4* dKO mouse heart during isoproterenol-induced sinus tachycardia. **B)** Optical map of action potentials (APs) shows ventricular activation in sinus tachycardia with regions of earliest activation (dark blue) at the exits of the right and left bundle branches. **C)** Pseudo-ECG shows frequent PVCs of morphology #1 in the same *Plxna3/a4* dKO mouse heart during isoproterenol stimulation, with **D)** earliest activation site at the lateral left ventricular (LV) wall. **E)** Pseudo-ECG shows frequent PVC and non-sustained VT of morphology #2 in the same mouse heart, with **F)** earliest activation site at the lateral right ventricular (RV) wall. **G)** Optical APs show temporal dissociation of atrial (magenta) and ventricular (blue) APs during optical mapping of PVC/VT #2.

### *Plxna3/a4* dKO mouse hearts display intrinsic adrenergic hypersensitivity *ex vivo*

The inducibility of VAs in *Plxna3/a4* dKO hearts with isoproterenol both *in vivo* and *ex vivo* in Langendorff led us to hypothesize that their VAs arise from intrinsic adrenergic hypersensitivity of the myocardium, rather than from overactivity of the sympathetic nervous system or increased circulating catecholamines. To test this hypothesis, we isolated hearts from *Plxna3/a4* dKO and WT mice and perfused them at constant pressure in an *ex vivo* Langendorff preparation, thus removing all autonomic nervous system and circulating catecholamine influence on cardiac physiology. In Langendorff, WT and *Plxna3/a4* dKO hearts had similar HR at baseline and similar peak HR with isoproterenol infusion (100 nM), and none of the hearts exhibited VAs at baseline during 20-minute pseudo-ECG recordings (Supplemental Figure 4). However, compared to WT, hearts from *Plxna3/a4* dKO mice demonstrated a significantly accelerated chronotropic response to isoproterenol (Figure 4A), with a shorter time to reach peak sinus rate (Figure 4B, Mann-Whitney test, p=0.0397).

**Figure 4.**
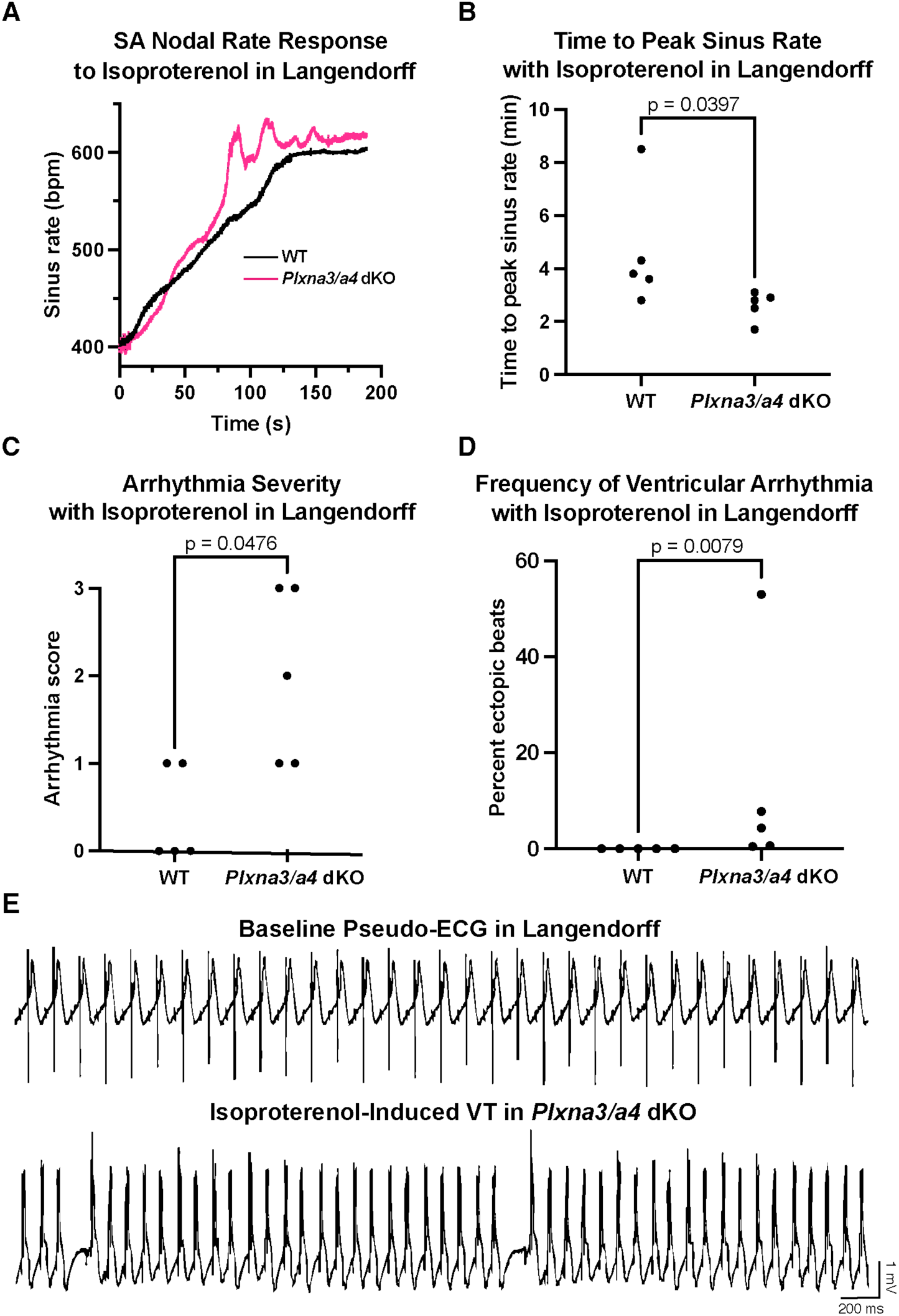
Hyperadrenergic electrophysiologic responses in Langendorff-perfused *Plxna3/a4* dKO hearts. **A)** Representative sinoatrial nodal rate response curves with isoproterenol perfusion (100 nM). **B)** Time to reach peak sinus rate after isoproterenol perfusion in *Plxna3/a4* dKO hearts versus WT. **C)** Isoproterenol-induced arrhythmia severity and **D)** frequency in Langendorff-perfused WT hearts versus *Plxna3/a4* dKO. All p-values are from Mann-Whitney tests and each dot in **(B-D)** represents one mouse. Arrhythmia severity scores: 0 = no PVCs, 1 = isolated PVCs, 2 = couplets or bigeminy, 3 = VT (3 or more consecutive ventricular ectopic beats). **E)** Representative pseudo-ECG tracings showing baseline sinus rhythm in a *Plxna3/a4* dKO heart (top) and sustained VT induced with isoproterenol (bottom).

Isoproterenol also induced VAs in all *Plxna3/a4* dKO hearts in Langendorff, with greater arrhythmia severity (p=0.0476, Mann-Whitney test) and frequency (p=0.0079, Mann-Whitney test) of ventricular ectopy, when compared with WT hearts (Figure 4C-D). In 2 of 8 *Plxna3/a4* dKO hearts, sustained VT was seen at peak isoproterenol effect, lasting several minutes (Figure 4E). These *ex vivo* findings suggest that the arrhythmic susceptibility in *Plxna3/a4* dKO mice is at the post-synaptic level, i.e. due to intrinsic hyperadrenergic response of the myocardium.

### *Plxna4* KO mice exhibit adrenergically driven ventricular arrhythmias due to increased post-synaptic β-adrenergic receptor density in ventricular membranes

Given the challenge of breeding *Plxna3/a4* double KO mice, we considered whether *Plxna4* single KO mice would also display denervation supersensitivity, since previous reports have shown that Plexin-A4 transduces the majority of Sema3a signaling in developing sympathetic nerves^25^, and *Plxna4* KO mice are also known to have decreased cardiac adrenergic innervation^26^. We performed 10-minute baseline ECG recordings in *Plxna4* KO mice and WT controls during isoflurane inhalation, which showed no baseline arrhythmias (data not shown), followed by 20-minute recordings after administration of isoproterenol (1 μg/g, i.p.). *Plxna4* KO mice displayed significantly higher frequency of VAs (Figure 5A, one-tailed Mann-Whitney test, p=0.0143) compared to WT controls. We confirmed the ventricular origin of these arrhythmias by optically mapping action potentials during PVCs and VT in two representative *Plxna4* KO hearts (Figure 5B).

**Figure 5.**
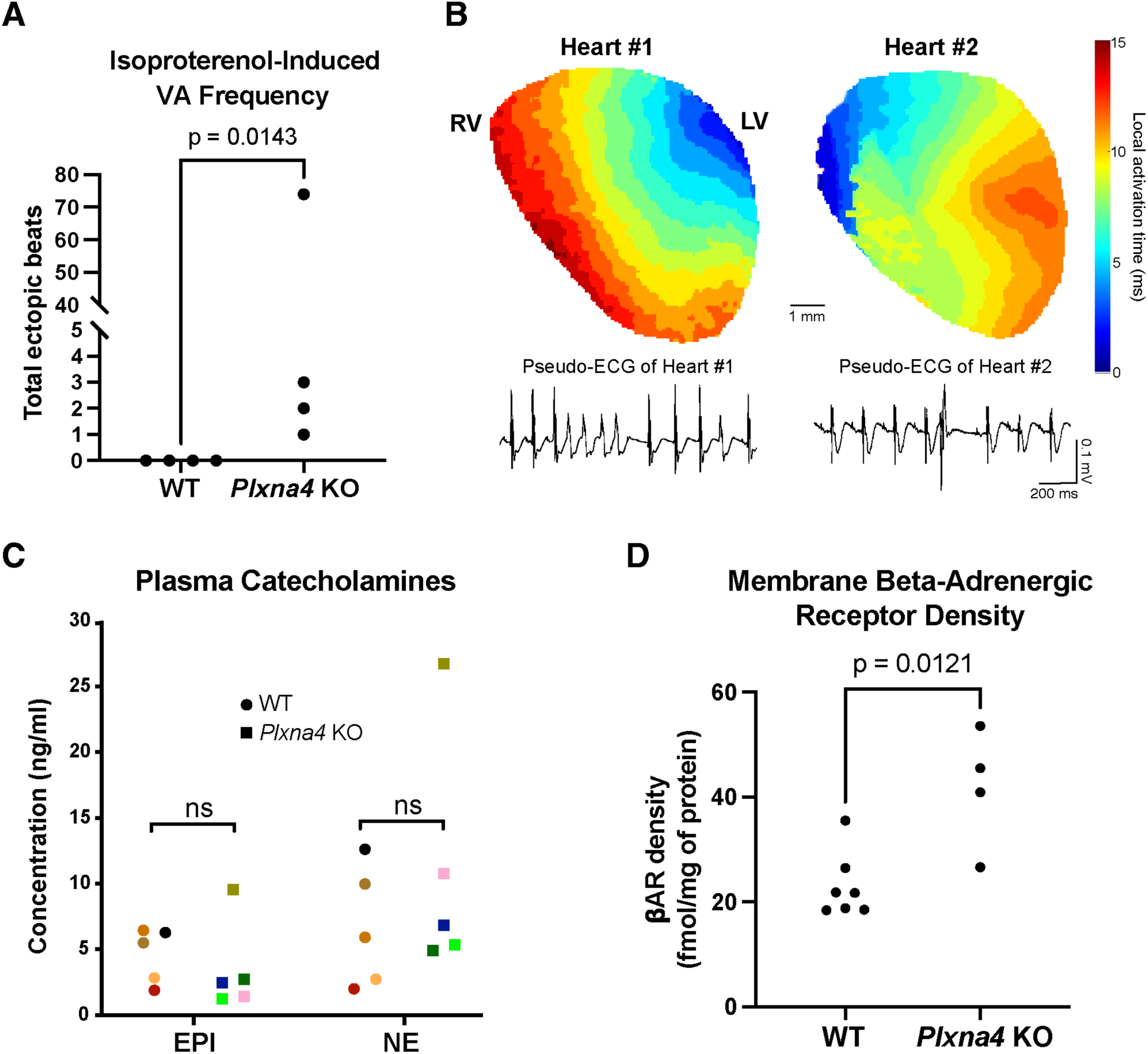
*Plxna4* KO mice have adrenergically driven ventricular arrhythmias and increased density of beta-adrenergic receptors in ventricular membranes. **A**) ECG recordings under anesthesia with isoflurane reveal that *Plxna4* KO mice have significantly increased isoproterenol-inducible ventricular arrhythmia (VA) frequency as measured by total ventricular ectopic beats during a 20-minute ECG recording (one-tailed Mann-Whitney test, each dot represents one mouse). **B)** Optical mapping of action potentials during VAs in two representative *Plxna4* KO mouse hearts show ventricular origins of the ectopic beats. **C)** Plasma catecholamine concentrations determined by ELISA were not significantly (ns) different in *Plxna4* KO mice compared to WT controls (Mann-Whitney tests, p>0.05). Data points with the same color are from the same mouse. **D)** Radioligand binding using the βAR antagonist cyanopindolol (125I-CYP) shows significantly higher βAR density in ventricular membranes of *Plxna4* KO mice when compared to WT hearts (Mann-Whitney test, each dot represents one mouse). EPI = epinephrine, NE = norepinephrine.

To determine the mechanism underlying the hyperadrenergic phenotype in *Plxna4* KO hearts, we measured circulating catecholamines in *Plxna4* KO mice by collecting plasma and quantifying epinephrine and norepinephrine levels with enzyme-linked immunosorbent assay (ELISA). There was no significant difference in plasma epinephrine and norepinephrine concentrations in *Plxna4* KO mice compared to WT controls (Figure 5C, Mann-Whitney test, p>0.05). We then extracted membranes from fresh frozen ventricular tissue of *Plxna4* KO mice and performed saturation binding assays using the radiolabeled beta-adrenergic receptor (βAR) antagonist [125I]-cyanopindolol. These assays showed an approximately two-fold increase in total βAR density (Bmax) in ventricular membranes from *Plxna4* KO mice compared to WT controls (Figure 5D, Mann-Whitney test, p = 0.0121). These findings support our contention that the pro-arrhythmic physiology following loss of plexin-dependent cardiac innervation is due to adrenergic hypersensitivity of the myocardium secondary to an increase in density of post-synaptic βARs.

### Human *PLXNA4* variants are associated with cardiac arrhythmias

To investigate the clinical relevance of our findings, we performed a phenome-wide association study (PheWAS) between phenotypes containing the word “arrhythmia” and the gene “*PLXNA4*” via the AstraZeneca PheWAS portal (https://azPheWAS.com/). This analysis identified 13 rare, single-nucleotide variants with nominally significant association p-values of <0.01 with arrhythmias (Figure 6A). One of these variants was a synonymous coding-region variant associated with two arrhythmia phenotype codes, with “re-entry ventricular arrhythmia” being the most significantly associated phenotype (Figure 6B), with a large effect size (odds ratio = 141). The p-values for individual variants did not reach the PheWAS significance threshold of p < 4.6e-6 (false discovery rate < 0.1)^32^. However, these associations found among different ancestries suggests that the *PLXNA4* gene may play a role in human arrhythmogenesis. The full variant list with detailed case-control comparisons and confidence intervals can be found in Supplemental Table 1.

**Figure 6.**
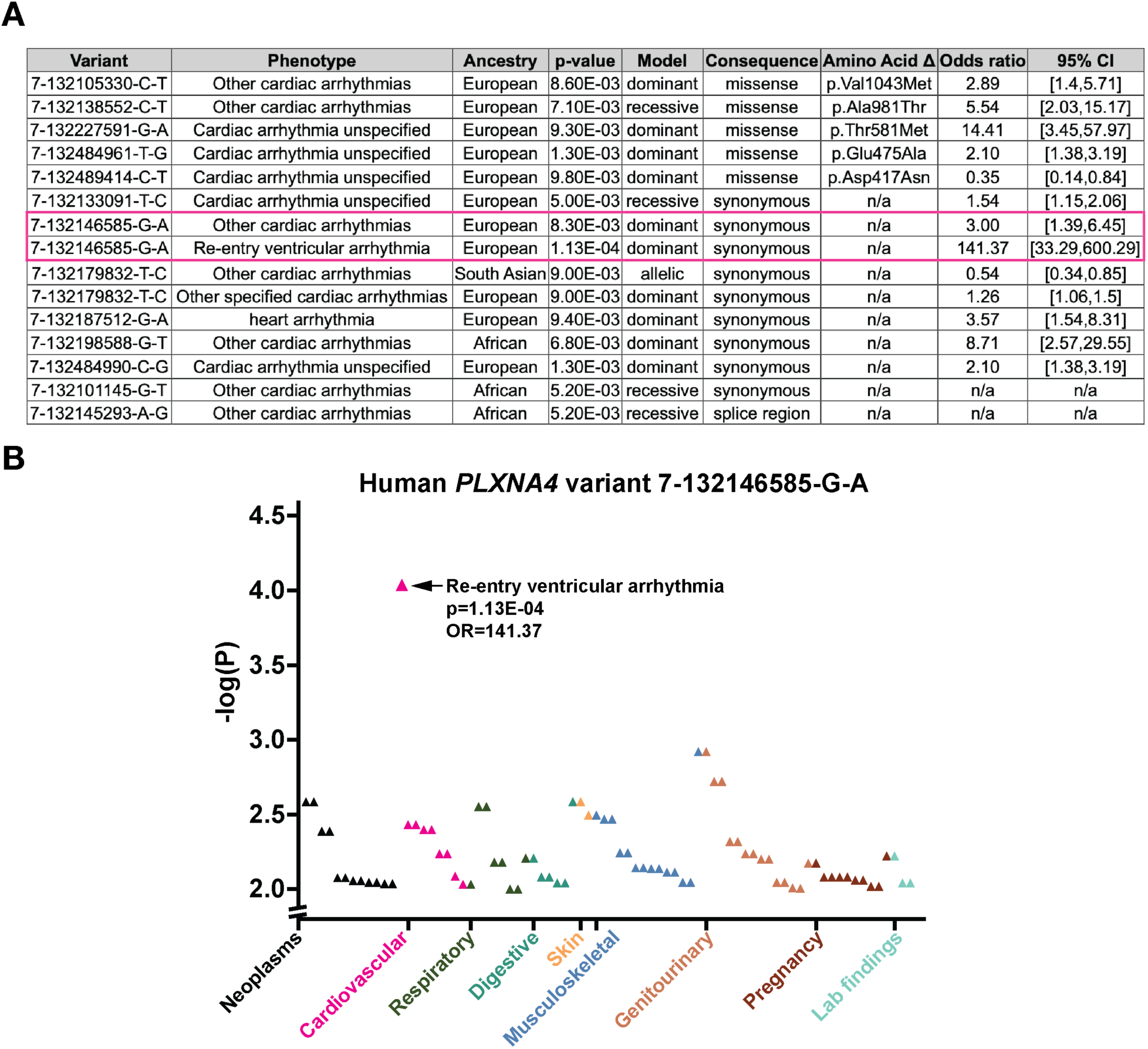
Association between rare *PLXNA4* variants and human arrhythmia phenotypes. Survey of the UK Biobank revealed **A)** 13 rare coding and splice single-nucleotide variants with nominally significant (p<0.01) associations with arrhythmia phenotypes, across several different ancestry groups and with several different inheritance patterns (models). One of these variants (highlighted in pink) was a synonymous variant associated with two arrhythmia phenotype codes. This variant was examined more closely in **B)** the Manhattan plot depicting the negative log-transformed P-values (y-axis) for its association with all phenotypes (x-axis), organized by disease categories. The most significantly associated phenotype for this variant was “Re-entry ventricular arrhythmia” (nominal p = 1.13e-4). OR = odds ratio. CI = confidence interval.

## DISCUSSION

In this study, we determined that loss of plexin-mediated cardiac innervation results in VAs driven by myocardial adrenergic hypersensitivity (Central Illustration). We demonstrated that this hyperadrenergic physiology is due to increased βAR density in cardiac membranes. This phenomenon is known as denervation supersensitivity, and it has previously been shown to be important in arrhythmogenic heart diseases such as MI and heart failure^13,33,34^, where there is injury to myocardium and degeneration of adrenergic axons. The precise molecular mechanisms of denervation supersensitivity are poorly understood. Our study highlights the importance of neuronal patterning mechanisms in regulating the adrenergic responsiveness of the myocardium. Recently, there has been increasing interest in the axon-guidance role of semaphorin-plexin signaling in arrhythmogenic heart diseases such as MI and heart failure^35,36^. The plexin ligand, Sema3A, has been studied for its protective effects after MI in animal models^22,24^ and for its potential as a biomarker for predicting risk of ventricular fibrillation after acute MI in humans^37^. These studies presume that the antiarrhythmic effect of Sema3A arises from its function in repelling sympathetic nerve sprouts after myocardial injury and thus decreasing adrenergic input, but this presumption has not yet been supported experimentally. As our data shows, a decrease in sympathetic innervation resulting from developmental loss of the Sema3A receptors, Plexin-A3 and –A4, is arrhythmogenic due to increased myocardial βAR density. Thus, sympathetic control over myocardial electrophysiology may vary widely in different disease states, and how nerve patterning determines arrhythmia susceptibility is far more complicated than a simple question of nerve quantity on the heart. The *Plxna3/a4* dKO mouse is a viable model that can be used to probe the specific role of cardiac nerve patterning in a variety of acquired heart diseases such as MI.

The observation that *Plxna3/a4* dKO mice exhibit spontaneous arrhythmias of ventricular origin, in the absence of cardiac structural abnormalities or intentional ion channel mutations, is particularly important. While atrial innervation is mostly parasympathetic, ventricular innervation is predominantly sympathetic^38,39^, and this may be why loss of plexin-mediated sympathetic innervation specifically leads to VAs. Compared with humans, mice are far less prone to arrhythmias in general due to numerous differences in their cardiac electrophysiology^40–42^. Even in humans, VAs are rare in the absence of myocardial injury or ion channelopathy^43^, and most channelopathies display incomplete penetrance of the arrhythmia phenotype^44^. Our finding that adrenergic stimulation without structural heart disease can cause sustained VT in *Plxna3/a4* dKO mouse hearts suggests that its physiology represents a unique platform for future studies of novel ion channel changes and specific adrenergic pathways driving VAs.

## LIMITATIONS

Several limitations of our study should be acknowledged. First, while we demonstrate a clear link between hypoinnervation and cardiac arrhythmogenesis, the precise molecular mechanisms by which deficiency of Plexin-A3 and Plexin-A4 leads to VAs requires further investigation, and we anticipate doing this by first using high-throughput screens of gene expression in *Plxna3/a4* dKO mouse hearts. Second, we identified several rare variants in human *PLXNA4* that were associated with arrhythmia phenotypes at nominally significant p-values of < 0.05. These phenotypes are based solely on ICD-9 diagnosis codes in the medical record, and further information on clinical diagnostics was not available. Thus the “Re-entry ventricular arrhythmia” phenotype could not be probed further in terms of actual arrhythmogenic mechanism.

Based on our experimental evidence thus far, *PLXNA4* variants would be anticipated to play a disease-modifying rather than a disease-causing role in human arrhythmias. While disease-causing arrhythmia genes, such as *KCNQ1*, frequently have association p-value that exceeded genome-wide significance, it is not surprising that PheWAS of *PLXNA4* as a likely disease modifier may not yield a significant p-value, when corrected for multiple-hypothesis testing. The absence of significant associations may also be due to negative selection of lethal arrhythmogenic variants and the relatively modest impact of even common variants on human arrhythmia. Though we recognize that PheWAS may be prone to false positive associations, the presence of multiple associated variants across different ancestries and variant types warrants future investigations in plexin-mediated cardiac innervation in human arrhythmias.

## CONCLUSION

In conclusion, our findings demonstrate that loss of plexin-mediated cardiac adrenergic innervation leads to ventricular arrhythmias through a mechanism involving post-synaptic βAR upregulation and enhanced adrenergic sensitivity, which may be relevant in human arrhythmogenesis. Adrenergic dysfunction is becoming an increasingly important area of investigation in arrhythmia biology, and neuromodulatory therapies have shown promise in the clinical management of VAs^45^. Thus, the *Plxna3/a4* dKO mouse is a novel tool for ongoing investigation into adrenergic mechanisms of arrhythmogenesis, which may identify new therapeutic targets for the prevention of sudden cardiac death.

## CLINICAL PERSPECTIVES

### Competency in Medical Knowledge

This is a pre-clinical study which offers novel insights into the mechanisms by which the adrenergic nervous system controls arrhythmia susceptibility. Neuromodulation targeting the adrenergic nervous system is an important part of complex arrhythmia management, therefore clinicians require knowledge of how neuromodulatory therapies exert their antiarrhythmic effects.

### Translational Outlook

The *Plxna3/a4* dKO mouse is a model of adrenergically driven VAs that may have relevance to human arrhythmogenesis. This model should be used in future basic and translational studies to elucidate pathways of neural patterning and adrenergic receptor signaling that may be novel antiarrhythmic targets.

## DATA AVAILABILITY

All data used in this study are available upon request to the corresponding author.

## AUTHOR CONTRIBUTIONS

C.Z., T.M., K.S., and H.A.R. designed the studies. C.Z. performed the animal surgeries, tissue collection, IF and tissue clearing, confocal imaging, telemetric ECG recordings, Langendorff experiments, optical mapping and data analyses. E.Y.L. performed cardiac tissue membrane extraction, radioligand binding assays, and mouse plasma collection. A.J. performed mouse plasma collection and catecholamine ELISA assays. T.M. and R.P. supplied mice and performed tissue section IF and imaging, as well as ECG recordings under anesthesia. J.J.W. assisted with the PheWAS. Y.C. performed echocardiography. C.Z. prepared the figures and wrote the manuscript with assistance from E.Y.L. and A.J. All coauthors contributed to the final version of the manuscript.

## ABBREVIATIONS

VA: ventricular arrhythmia
ECG: electrocardiography / electrocardiogram
dKO: double knockout
IF: immunofluorescence
TH: tyrosine hydroxylase
PVC: premature ventricular contraction
PAC: premature atrial contraction
VT: ventricular tachycardia
βAR: beta-adrenergic receptor
PheWAS: phenome-wide association study

## ACKNOWLEDGEMENTS

We are grateful to Dr. Kristina Bostrom and Dr. Conrad Hodgkinson for their thoughtful review and suggestions for the manuscript. This work was supported by an American Heart Association Career Development Award (24CDA1268188) to C.Z. and NIH grants: Stimulating Peripheral Activity to Relieve Conditions (OT2OD023848) and P01HL164311 to K.S., and R01HL056687 to H.A.R.

---

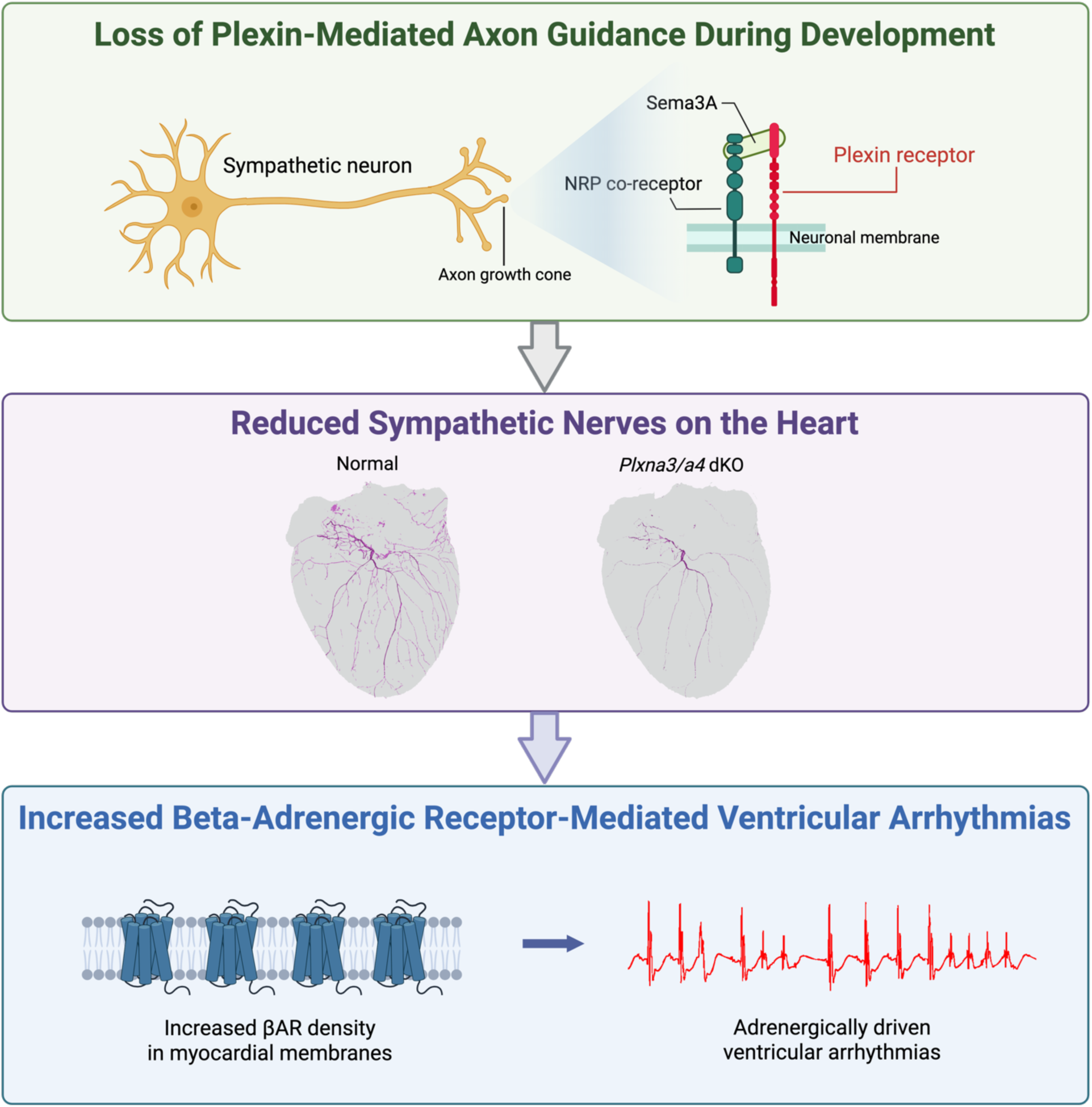
Central Illustration. NRP = neuropilin. dKO = double knockout. βAR = beta-adrenergic receptor.

