## Supplemental Materials for "Adrenergic Hypersensitivity Drives Ventricular Arrhythmias Following Loss of Plexin-Mediated Cardiac Innervation"

**Supplemental Appendix**

**Contents**

Supplemental Table 1. Detailed PheWAS results p. 2

Supplemental Figure 1. Masson's trichrome stain of *Plxna3/a4* dKO heart sections p. 4

Supplemental Figure 2. Atrial innervation and atrial arrhythmias in *Plxna3/a4* dKO versus WT mice p. 5

Supplemental Figure 3. Telemetric ECG parameters for *Plxna3/a4* dKO versus WT mice p. 6

Supplemental Figure 4. Langendorff pseudo-ECG parameters for *Plxna3/a4* dKO versus WT hearts p. 7

| Variant | Phenotype | Ancestry | p-value | Model | Consequence type | Transcript | cDNA change | Amino acid change | Exon rank | No. cases | No. AA cases | No. AB cases | No. BB cases | Case AAF | Proportion AB or BB cases | Proportion BB cases | No. controls | No. AA controls | No. AB controls | No. BB controls | Control AAF | Proportion AB or BB controls | Proportion BB controls | Odds ratio | Odds ratio LCI | Odds ratio UCI |
|---|---|---|---|---|---|---|---|---|---|---|---|---|---|---|---|---|---|---|---|---|---|---|---|---|---|---|
| 7-132489414-C-T | 41202#I499#Cardiac arrhythmia unspecified | European | 0.0098 | dominant | missense_variant | ENST00000321063 | c.1249G>A | p.Asp417Asn | 3/32 | 611 | 606 | 5 | 0 | 0.0041 | 0.00818331 | 0 | 245795 | 240084 | 5667 | 44 | 0.0117 | 0.02323481 | 0.00017901 | 0.3469 | 0.1438 | 0.8367 |
| 7-132484990-C-G | 41202#I499#Cardiac arrhythmia unspecified | European | 0.0013 | dominant | synonymous_variant | ENST00000378539 | c.1395G>C | p.Gly465Gly | 5/5 | 611 | 588 | 23 | 0 | 0.0188 | 0.03764321 | 0 | 245794 | 241304 | 4468 | 22 | 0.0092 | 0.01826733 | 8.9506E-05 | 2.1022 | 1.3845 | 3.1919 |
| 7-132484961-T-G | 41202#I499#Cardiac arrhythmia unspecified | European | 0.0013 | dominant | missense_variant | ENST00000378539 | c.1424A>C | p.Glu475Ala | 5/5 | 611 | 588 | 23 | 0 | 0.0188 | 0.03764321 | 0 | 245794 | 241304 | 4468 | 22 | 0.0092 | 0.01826733 | 8.9506E-05 | 2.1022 | 1.3845 | 3.1919 |
| 7-132227591-G-A | 41202#I499#Cardiac arrhythmia unspecified | European | 0.0093 | dominant | missense_variant | ENST00000321063 | c.1742C>T | p.Thr581Met | 7/32 | 611 | 609 | 2 | 0 | 0.0016 | 0.00327332 | 0 | 245801 | 245745 | 55 | 1 | 0.0001159 | 0.00022783 | 4.0683E-06 | 14.4115 | 3.509 | 59.1885 |
| 7-132198588-G-T | 131353#Source of report of I49 (Other cardiac arrhythmias) | African | 0.0068 | dominant | synonymous_variant | ENST00000321063 | c.2635C>A | p.Arg879Arg | 13/32 | 143 | 140 | 3 | 0 | 0.0105 | 0.02097902 | 0 | 8561 | 8540 | 21 | 0 | 0.0012 | 0.00245298 | 0 | 8.7143 | 2.5696 | 29.5527 |
| 7-132198552-C-T | 41202#I49#Other cardiac arrhythmias | European | 0.0071 | recessive | missense_variant | ENST00000321063 | c.2671G>A | p.Ala891Thr | 13/32 | 2520 | 2436 | 80 | 4 | 0.0175 | 0.03333333 | 0.0015873 | 258093 | 249832 | 8187 | 74 | 0.0161 | 0.03200784 | 0.00028672 | 5.5433 | 2.0252 | 15.1727 |
| 7-132187512-G-A | 20002#1077#heart arrhythmia | European | 0.0094 | dominant | synonymous_variant | ENST00000321063 | c.2952C>T | p.Asn984Asn | 15/32 | 7011 | 7005 | 6 | 0 | 0.0004279 | 0.0008558 | 0 | 221095 | 221042 | 53 | 0 | 0.0001199 | 0.00023972 | 0 | 3.5723 | 1.5351 | 8.3126 |
| 7-132185330-C-T | 40002#I49#Other cardiac arrhythmias | European | 0.0086 | dominant | missense_variant | ENST00000321063 | c.3127G>A | p.Val1043Met | 16/32 | 156 | 148 | 8 | 0 | 0.0256 | 0.05128205 | 0 | 247455 | 242915 | 4514 | 26 | 0.0092 | 0.01834677 | 0.00010507 | 2.8922 | 1.419 | 5.8948 |

| Variant | Phenotype | Ancestry | p-value | Model | Consequence type | Transcript | cDNA change | Amino acid change | Exon rank | No. cases | No. AA cases | No. AB cases | No. BB cases | Case AAF | Proportion AB or BB cases | Proportion BB cases | No. controls | No. AA controls | No. AB controls | No. BB controls | Control AAF | Proportion AB or BB controls | Proportion BB controls | Odds ratio | Odds ratio LCI | Odds ratio UCI |
|---|---|---|---|---|---|---|---|---|---|---|---|---|---|---|---|---|---|---|---|---|---|---|---|---|---|---|
| 7-132179832-T-C | Union#I498#Other specified cardiac arrhythmias | European | 0.009 | dominant | synonymous_variant | ENST00000321063 | c.3729A>G | p.Ala1243Ala | 20/32 | 1081 | 144 | 516 | 421 | 0.6281 | 0.86679001 | 0.38945421 | 140724 | 22824 | 67631 | 50269 | 0.5975 | 0.83781018 | 0.35721696 | 1.2597 | 1.0564 | 1.5021 |
| 7-132164145-G-T | 41202#I49#Other cardiac arrhythmias | African | 0.0052 | recessive | synonymous_variant | ENST00000321063 | c.4497C>A | p.Thr1499Thr | 24/32 | 37 | 36 | 0 | 1 | 0.027 | 0.02702703 | 0.02702703 | 7018 | 6887 | 131 | 0 | 0.0093 | 0.01866629 | 0 | - | - | - |
| 7-132146585-G-A | Union#I49#Other cardiac arrhythmias | European | 0.0083 | dominant | synonymous_variant | ENST00000321063 | c.4980C>T | p.His1660His | 28/32 | 16947 | 16938 | 9 | 0 | 0.0002655 | 0.00053107 | 0 | 135431 | 135407 | 24 | 0 | 0.00008861 | 0.00017721 | 0 | 2.9979 | 1.3932 | 6.4505 |
| 7-132145293-A-G | 41202#I49#Other cardiac arrhythmias | African | 0.0052 | recessive | splice_region_variant | ENST00000321063 | c.5056-5T>C | NA | 28/31 | 37 | 36 | 0 | 1 | 0.027 | 0.02702703 | 0.02702703 | 7018 | 6878 | 140 | 0 | 0.01 | 0.0199487 | 0 | - | - | - |
| 7-132133091-T-C | Union#I499#Cardiac arrhythmia unspecified | European | 0.005 | recessive | synonymous_variant | ENST00000321063 | c.5547A>G | p.Ala1849Ala | 31/32 | 3300 | 2699 | 553 | 48 | 0.0983 | 0.18212121 | 0.01454545 | 144155 | 117110 | 25678 | 1367 | 0.0985 | 0.18761056 | 0.00948285 | 1.5417 | 1.1537 | 2.0603 |

**Supplemental Table 1. Detailed phenome-wide association study results.**

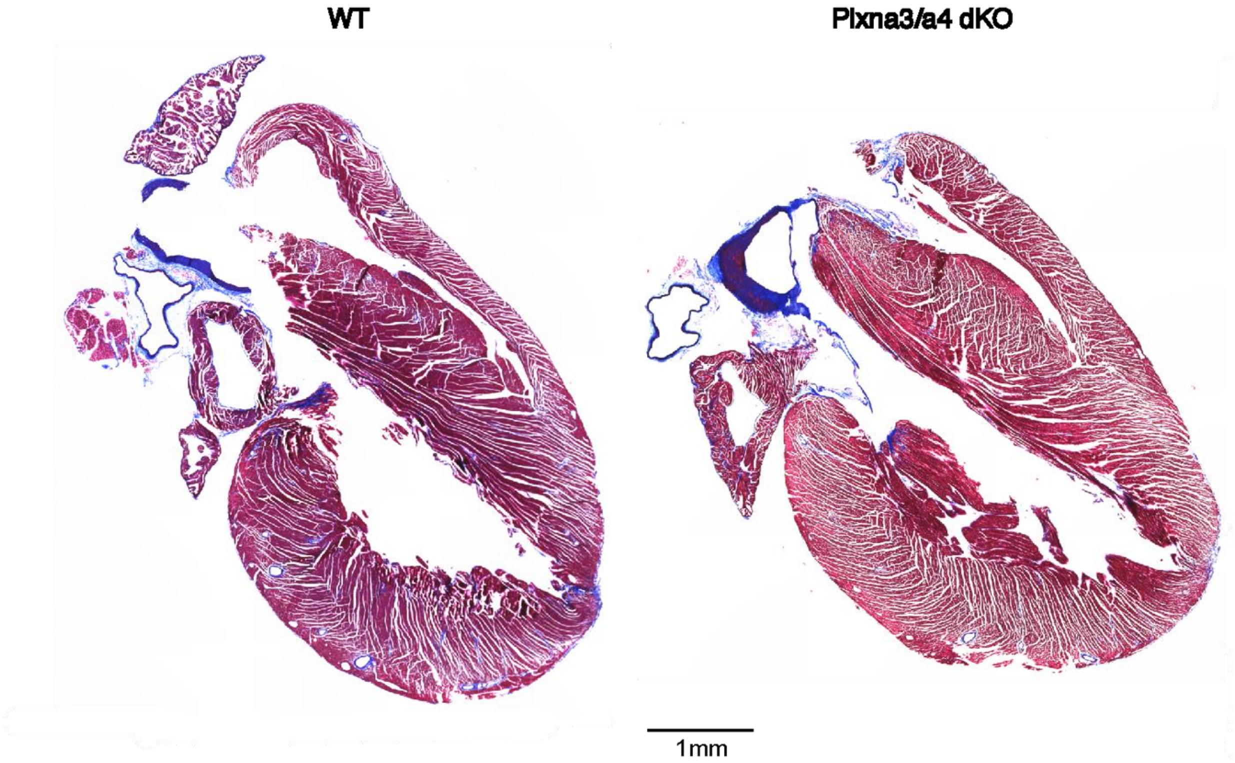


**Supplementary Figure 1. Representative sections from *Plxna3/a4* KO hearts show no signs of fibrosis when stained with Masson's trichrome**.

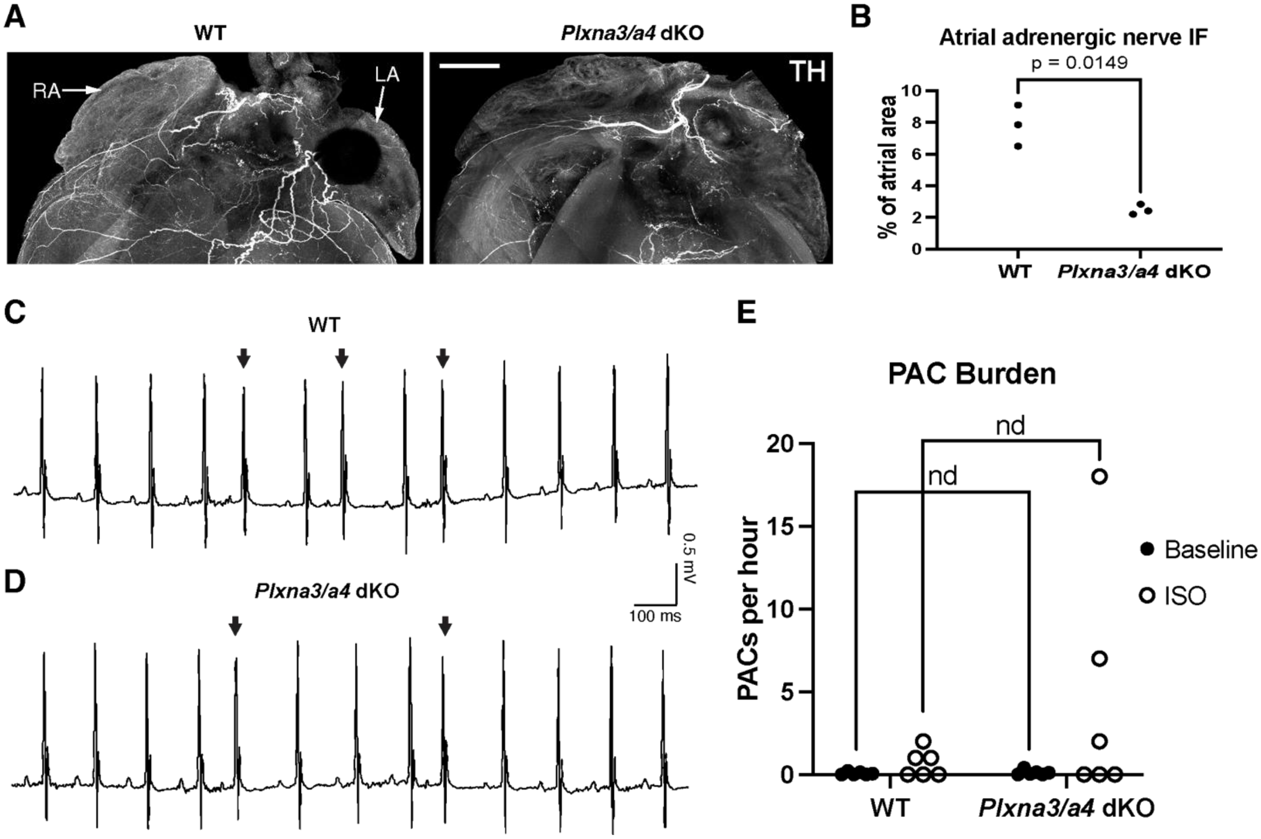


**Supplementary Figure 2. Atrial innervation and atrial arrhythmias in *Plxna3/a4* dKO versus WT mice**. **A-B)** Sympathetic nerve density was significantly reduced in atria of *Plxna3/a4* dKO mice compared to WT mice (Welch's t-test, each dot represents one mouse). Very rare spontaneous premature atrial contractions (PACs) were seen in both **(C)** WT and **(D)** *Plxna3/a4* dKO mice during conscious telemetric ECG recordings. **E)** There was no difference in PAC burden at baseline or after isoproterenol administration (multiple unpaired t-tests with two-stage step-up method of false discovery rate correction for multiple comparisons, $p > 0.1$ for both comparisons, $n = 6$ mice per group). TH = tyrosine hydroxylase. nd = not a discovery, ISO = isoproterenol, WT = wildtype, KO = knockout. Scale bar in **(A)** = 200μm.

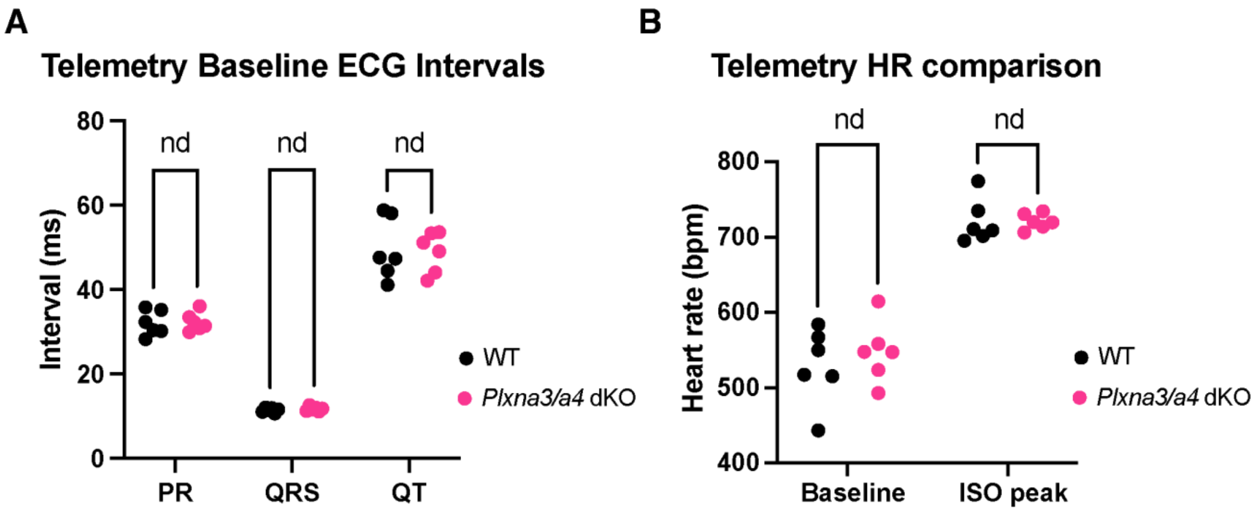


**Supplementary Figure 3. Telemetric ECG parameters for *Plxna3/a4* dKO versus WT mice**. There was no difference in baseline ECG intervals, baseline resting HR, or peak HR after isoproterenol administration (multiple unpaired t-tests with two-stage step-up method of false discovery rate correction for multiple comparisons, $p > 0.3$ for all comparisons, $n = 6$ mice per group). HR = heart rate, nd = not a discovery, ISO = isoproterenol, WT = wildtype, KO = knockout.

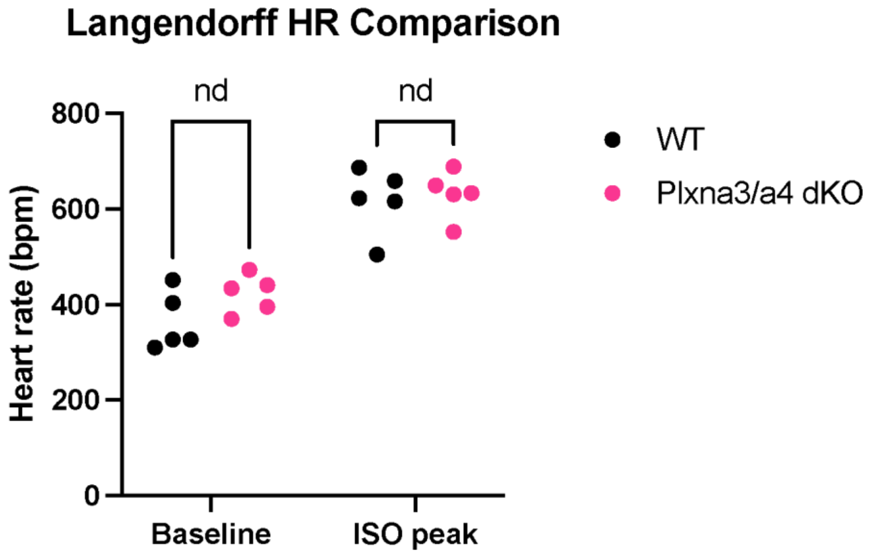


**Supplementary Figure 4. Langendorff pseudo-ECG parameters for *Plxna3/a4* dKO versus WT hearts**. There was no difference in baseline HR or peak HR after isoproterenol administration (multiple unpaired t-tests with two-stage step-up method of false discovery rate correction for multiple comparisons, $p > 0.1$ for both comparisons, $n = 6$ mice per group). HR = heart rate, nd = not a discovery, ISO = isoproterenol, WT = wildtype, KO = knockout.
